# The transcription factor SpiB regulates Fibroblastic Reticular Cell network and CD8^+^ T cell responses in lymph nodes

**DOI:** 10.1101/2023.10.12.562124

**Authors:** Harry L Horsnell, Wang HJ Cao, Gabrielle T Belz, Scott N Mueller, Yannick O Alexandre

## Abstract

Fibroblastic Reticular Cells (FRCs) construct microanatomical niches that support lymph node homeostasis and coordination of immune responses. Transcription factors regulating the functionality of FRCs remain poorly understood. Here we investigate the role of the transcription factor SpiB that is expressed in lymph node FRCs. Conditional ablation of SpiB in FRCs impaired the FRC network in the T cell zone of lymph nodes, leading to reduced numbers of FRCs and altered homeostatic functions including reduced CCL21 and interleukin-7 expression. The size and cellularity of lymph nodes remained intact in the absence of SpiB but the space between the reticular network increased, indicating that although FRCs were reduced in number they stretched to maintain network integrity. Following virus infection, antiviral CD8^+^ T cell responses were impaired, suggesting a role for SpiB expression in FRCs in orchestrating immune responses. Together, our findings reveal a new role for SpiB as a critical regulator of FRC functions and immunity in lymph nodes.

## Introduction

Lymph nodes (LNs) are secondary lymphoid organs that form a network of tissues designed as a filtration and surveillance system. The microarchitecture of LNs is organised into distinct compartments with the primary goal of capturing and presenting antigens from peripheral tissues to cells of the immune system, generating immune responses^1^. As such, dendritic cells present antigens to T cells for priming in the T cell zone, whereas B cell responses and the formation of germinal centres occurs in B cell follicles. The LN subcapsular sinus operates as a first line of defence with a layer of specialised subcapsular sinus macrophages that sample afferent lymph and capture pathogens and antigens that drain from the tissues^2^. The medulla contains blood vessels and medullary lymphatics that serve as an exit route for lymphocytes back into the blood circulation. Several types of non-haematopoietic stromal cells support the lymphoid architecture and function by constructing networks and defining compartments. CD31^+^ lymphatic and blood endothelial cells build the vasculature of LNs required for the entry and exit of immune cells. Contractile pericytes expressing the adhesion molecule CD146 are also found in association with blood vessels. Fibroblastic Reticular Cells (FRCs) are the most prominent stromal cell population in LNs that form an interconnected cellular network that supports immune cell migration^3^. In addition, FRCs create a conduit system composed of extracellular matrix components and reticular fibers that facilitates the transport of lymph-derived antigens and signalling molecules, assisting in the induction of immune responses. Inflammation transcriptionally reprograms FRCs towards immune related pathways that further support ongoing immune responses^4–6^.

FRCs comprise several subsets based on their intranodal location, markers, and functions, that all express podoplanin (PDPN). This heterogeneity in FRCs creates anatomical niches that support the homeostasis and support of immune responses in LNs^7^. Within the subcapsular sinus, marginal reticular cells (MRCs) and lymphatic endothelial cells form a unique niche for subcapsular macrophage development and homeostasis^8–10^. B cell zone reticular cells include follicular dendritic cells (FDCs) and other CXCL13-expressing reticular cells that define this compartment and support humoral responses^11^. Within the T cell zone, reticular cells (TRCs) express the chemokines CCL19 and CCL21 that promote the attraction and retention of T cells and DC that express CCR7 and produce cytokines such as interleukin (IL)-7 that are critical for T cell survival ^12,13^. Additionally, MRCs, BRCs and TRCs are all characterised by high expression of the bone marrow stromal cell antigen-1, or CD157. Conversely, in the medulla CD157^low^ FRCs (medRC) were recently shown to support plasma cells by providing IL-6 ^7,14^. Finally, adventitial reticular cells (ARCs) that express CD34 and Ly6C form a specific niche by surrounding blood vessels and may function as precursors of adult FRCs^15,16^.

LN FRCs develop from fibroblast activation protein-α^+^ lymphoid tissue organiser cells (LTos) of mesenchymal origin^17^. Their development requires sequential differentiation and maturation steps that are not fully understood. Several pathways such as the lymphotoxin-β receptor signalling, NF-κB or effectors of Hippo signalling, YAP and TAZ were shown to instruct the maturation of FRCs from FRC precursor cells^18–21^. Deficiency in these pathways results in a reduction in the cellularity of FRCs and these immature FRCs harbor lower expression of the homeostatic chemokines CCL19, CCL21 and CXCL13 as well as reduction in IL-7. Interestingly, immature FRCs still produce reticular fibres and a functional conduit network in LNs, but the overall size and immune cell cellularity is decreased^18,19^. These defects in the FRC network also resulted in impaired CD8^+^ T cell responses during viral infections, indicating that the generation of optimal immune responses require a functional and healthy FRC network. However, it is not clear how the FRC network impacted antiviral immunity because in those studies the position and recruitment of immune cells in LNs were also perturbed^18,19^.

We previously identified the transcription factor SpiB as a regulator of FRC maturation in the spleen^16,22^. Here we investigated the role of SpiB in LN homeostasis and immune responses. We found that SpiB expression is conserved in LN FRCs and conditional ablation of SpiB expression impacted the cellularity and functionality of the FRC network in the T cell zone at steady state. Deletion of SpiB in FRCs did not affect LN tissue size or homeostasis of immune cells but impacted T cell priming following viral infection. These data indicate conserved function of the transcription factor SpiB in lymphoid organ FRCs for the induction of pathogen-specific T cell responses.

## Results

### The transcription factor SpiB is expressed by lymph node FRC

We previously identified a role for SpiB in FRCs in the spleen^16^. We first sought to determine if SpiB expression was conserved in LNs. Integration of scRNA-seq datasets analysing FRCs from both spleen and LNs revealed expression of *Spib* mRNA in FRCs from LN B and T cell zones as well as medullary reticular cells (medRCs) but low to no expression in LN ARCs or pericytes (Supp Fig. 1A). B cell follicle FRCs had higher expression of *Spib* compared to TRCs. We then used SpiB-tdTomato reporter mice to examine expression in LN stromal cells by flow cytometry and observed that FRCs and LECs had the highest levels of tdTomato whilst BEC and pericytes had low expression (Fig. 1A-B). We then used a new gating strategy to further define FRC subsets in LNs. We identified CD21/35^+^ FDC, MadCAM1^+^ MRCs, CD157^+^ TRC, Ly6C^+^CD157^-^ ARCs and Ly6C^-^CD157^-^ medRCs (Fig. 1C)^14,16,23^. Flow cytometry analysis revealed that FDCs and MRCs had the highest expression of tdTomato followed by TRC, whilst ARC and medRC had low levels of tdTomato (Fig. 1D).

**Figure 1:**
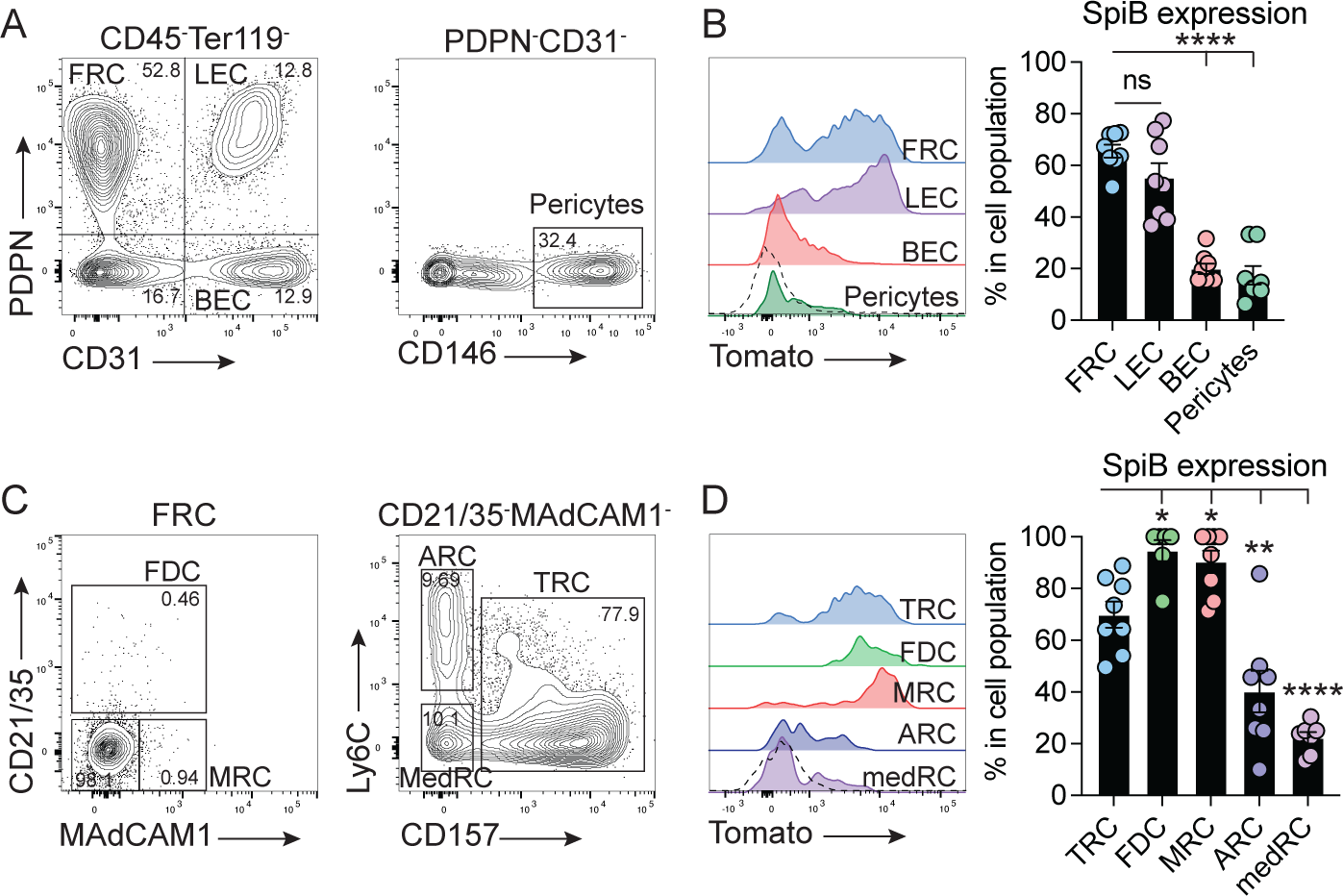
LN FRCs express the transcription factor SpiB. **(A)** Gating strategy to identify lymph node (LN) stromal cells by flow cytometry. Hematopoietic cells were excluded, and stromal cells were identified with the markers PDPN, CD31 and CD146. FRC: Fibroblastic Reticular Cells; LEC: Lymphatic endothelial cells; BEC: Blood endothelial cells. **(B)** Flow cytometry of LN stromal cell subsets in SpiB-TdTomato mice. Representative histograms of SpiB expression (left) and pooled data (means ± SEM) from eight mice combined from three experiments. Dotted line represents baseline fluorescence from WT control. **(C)** Gating strategy to identify lymph node FRC subsets by flow cytometry. FDC: Follicular Dendritic Cells; MRC: Marginal Reticular cells; ARC: Adventitial Reticular Cells; MedRC: Medullary Reticular Cells; TRC: T zone Reticular Cells. **(D)** Flow cytometry of LN stromal cell subsets in SpiB-TdTomato mice. Representative histograms of SpiB expression (left) and pooled data (means ± SEM) from eight mice combined from three experiments. Dotted line represents baseline fluorescence from WT control. *P < 0.05, **P < 0.01, and ****P < 0.0001, ns, non-significant, by ANOVA with Tukey’s multiple comparisons test (B and D).

### The transcription factor SpiB supports FRC homeostasis

To investigate a role for SpiB in LN FRCs, we used Ccl19-Cre/Spib^flox/flox^ mice (SpiB^ΔCCL19^) as previously described^16^ and confirmed the absence of *Spib* in sorted populations of TRCs, ARCs and medRCs by qPCR (Supp Fig. 1B). We observed a significant reduction in the numbers of FRCs, LECs and BECs but not of pericytes in the LN of SpiB^ΔCCL19^ mice (Fig. 2A). The reduction in LECs and BECs may reflect a bystander effect because endothelial cells are not targeted in CCL19-Cre mice^19,24^. Amongst FRC subsets, TRCs and MRCs were reduced in cellularity in SpiB^ΔCCL19^ mice but FDCs, ARCs and medRCs were not changed (Fig. 2B). Thus, expression of SpiB was required for maintenance of the stromal cell networks that support LNs in defined areas of LNs.

**Figure 2:**
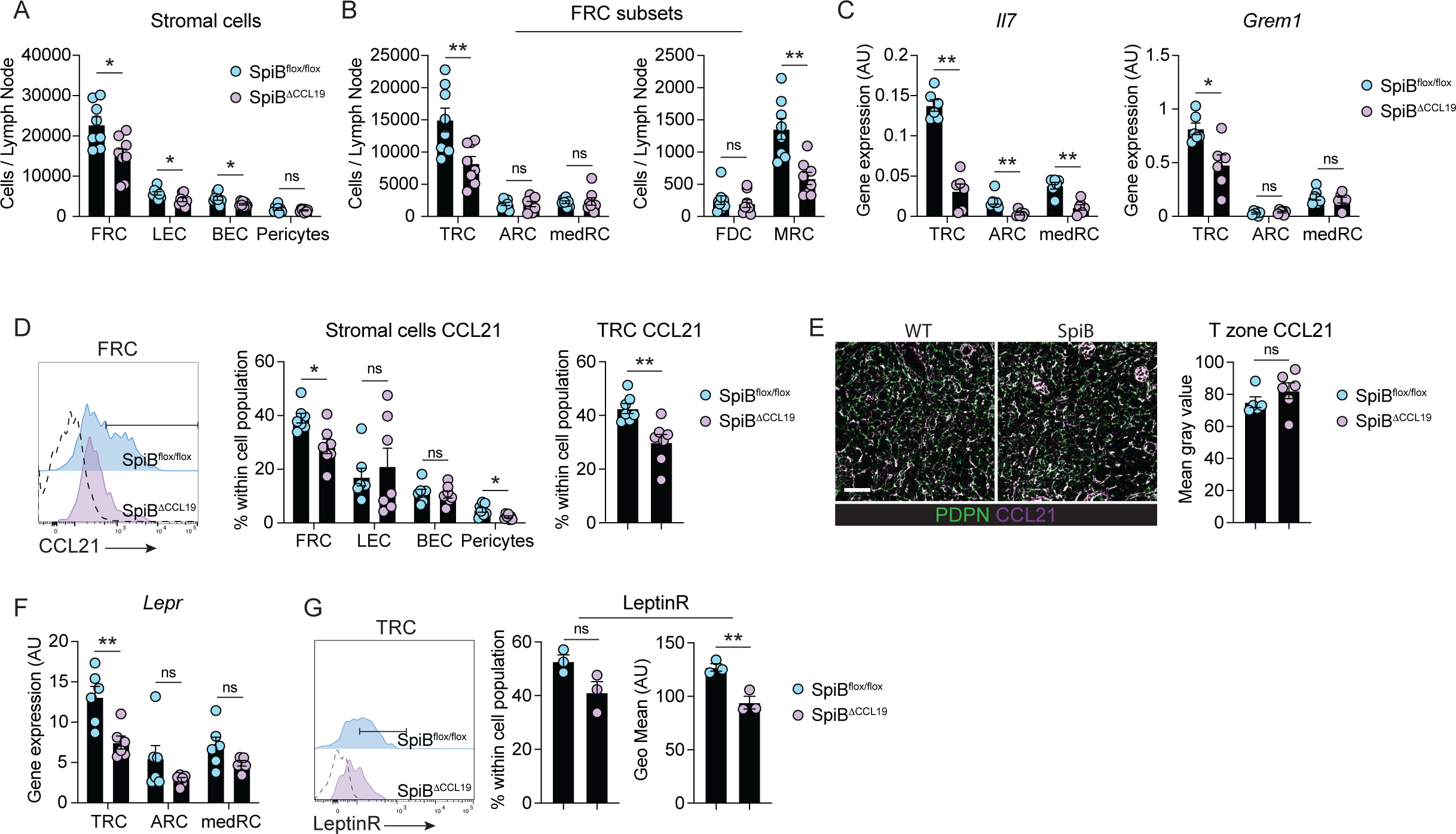
The transcription factor SpiB controls the T zone reticular cell network and functionality. **(A-B)** Enumeration of lymph node stromal cells and FRC subsets from control SpiB^Flox/Flox^ and SpiB^ΔCCL19^ mice by flow cytometry. Graphs show pooled data (means ± SEM) from two independent experiments with 8 mice per group. **(C)** Analysis of *Il7* and *Grem1* expression in lymph node sorted TRCs, ARC and medRC of control SpiB^Flox/Flox^ and SpiB^ΔCCL19^ mice by qPCR. n = 6 mice from 3 independent sorts. **(D)** Flow cytometry analysis and representative histograms (left) of intracellular CCL21 expression in FRCs from control SpiB^Flox/Flox^ and SpiB^ΔCCL19^. Fluorescence minus one staining is shown by the histogram with a dotted line and used to discriminate CCL21^+^ from CCL21^-^ cells. Percentage (right) of CCL21^+^ stromal cell subsets and TRCs in the LNs of control SpiB^Flox/Flox^ and SpiB^ΔCCL19^ mice. Graphs show pooled data (means ± SEM) from two independent experiments with 7 mice per group. **(E)** Skin draining lymph node sections from control SpiB^Flox/Flox^ and SpiB^ΔCCL19^ mice were stained for PDPN and CCL21 and analysed by confocal microscopy, and the area of CCL21 area was quantified in the T cell zone of LN. Graph shows pooled data (means ± SEM) from two independent experiments with 5-6 mice per group. Scale bar, 50 μm. **(F)** Analysis of *Lepr* expression in lymph node sorted TRCs, ARC and medRC of control SpiB^Flox/Flox^ and SpiB^ΔCCL19^ mice by qPCR. n = 6 mice from 3 independent sorts. **(G)** Flow cytometry analysis of LepR expression in TRCs from control SpiB^Flox/Flox^ and SpiB^ΔCCL19^ mice. Left: representative histograms of LepR staining with Fluorescence minus one staining shown as dotted line and used to discriminate LepR^+^ from LepR^-^ cells. Right: graphs show the percentage of LepR^+^ TRCs in LNs and the Mean Fluorescence Intensity (GeoMean) of LepR in TRCs. Data are representative of one experiment out of two with three mice per group. *p < 0.05, **p < 0.01, ****p < 0.0001, ns, non-significant, by unpaired two-tailed t test (A-B and G) and Mann-Whitney test (C-F).

We previously showed that in the absence of SpiB, splenic TRCs had increased expression of genes expressed by ARCs, notably the stem marker CD34, and decreased expression of mature TRC markers, including *Il7*, *Ccl19* and *Grem1*, which indicated a role for SpiB in supporting the differentiation of spleen TRCs from adventitial precursor cells^16^. We did not observe changes in *Ccl19* expression in SpiB-deficient LN TRCs (Supp Fig. 1C), however, expression of the homeostatic cytokine *Il7* was significantly reduced in FRC subsets and *Grem1 as well as Cd34* expression were reduced in TRCs (Fig. 2C and Supp Fig. 1C). This suggests that SpiB likely regulates some aspects of the differentiation of FRCs in LNs. We then asked if SpiB regulates the expression of other canonical FRC markers in FRCs, including the proteins PDPN, CD157, VCAM-1 and CD140a (PDGFRa). We observed small changes in expression of these markers in MRCs in the absence of SpiB and no differences amongst other LN FRC subsets except for CD140a that was also slightly reduced in LN TRCs (Supp Fig. 1D). In addition, SpiB^ΔCCL19^ FRCs demonstrated normal expression of the chemokines *Ccl2*, *Ccl7*, *Cxcl9*, *Cxcl10*, *Cxcl12*, *Cxcl13* and the alarmin *Il33*, except for a reduction of *Ccl7* expression in TRCs and *Cxcl13* in medRC (Supp Fig. 1E). We observed a significant reduction in intracellular CCL21 expression in SpiB^ΔCCL19^ FRCs, particularly within TRCs, but not in endothelial cells or pericytes (Fig. 2D). Histological examination showed no difference in CCL21 deposition in the T cell zone of SpiB^ΔCCL19^ mice (Fig. 2E), possibly due to accumulation of the chemokine on the reticular network via heparan sulphate binding^25^. LN FRCs that surround HEV and LEC also predominantly express the leptin receptor (LepR), which might promote FRC survival and functions^26^. We found that *Lepr* was reduced in TRCs in the absence of SpiB and confirmed the decrease of LepR expression by flow cytometry in SpiB-deficient TRCs (Fig 2. F-G). Overall, these data show that SpiB regulates the maintenance of TRCs and MRCs in LNs and regulates discreet components of FRC function.

### Normal lymph node architecture in mice lacking SpiB in FRCs

Because FRCs play crucial roles in regulating LN homeostasis, we investigated if the reduction in numbers of TRCs and altered functionality impacted immune cell cellularity and the LN architecture. The loss of SpiB in FRCs resulted in a small but non-significant decrease in total LN cellularity, composed of small but non-significant changes in the numbers of B cells, CD4^+^ and CD8^+^ T cells (Fig. 3A). Other immune cells including NK and NK-T cells, monocytes, neutrophils and dendritic cell subsets, apart from plasmacytoid dendritic cells, were not altered in the LNs of SpiB^ΔCCL19^ mice (Fig. 3A). The gross architecture and total surface area of LNs remained unchanged in SpiB^ΔCCL19^ mice (Fig. 3B-C). The absence of SpiB expression in FRCs also did not affect the organisation or size of the LN T cell zone, B cell follicles or medulla (Fig. 3C). However, a closer examination of the PDPN^+^ TRC network in the T cell zone revealed that the TRC network was less dense in SpiB^ΔCCL19^ mice (Fig. 3D). Quantification of the spacing between FRCs in the T cell zone network using gap analysis confirmed that the space between the reticular network fibers was increased in the absence of SpiB (Fig. 3E). The reduction in FRC and lymphocyte cellularity resulted in maintenance of the ratio of T cells to TRCs (Fig. 3F). Thus, increased spacing between TRCs in LNs from SpiB^ΔCCL19^ mice occurred in the absence of a reduction in tissue size, suggesting that the FRC network stretched to maintain T cell homeostasis.

**Figure 3:**
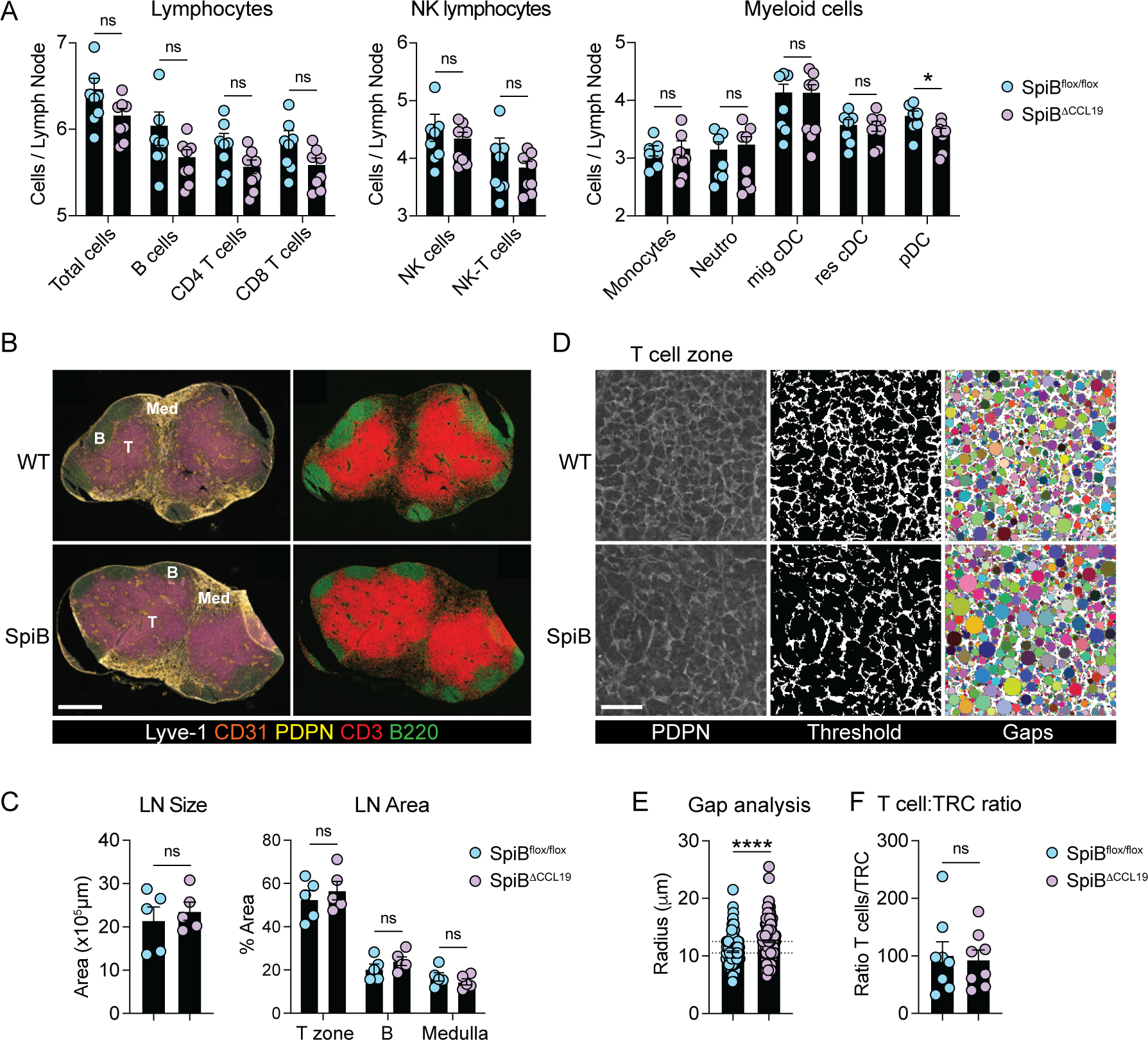
SpiB deletion in FRC does not impact immune cell homeostasis or LN architecture. **(A)** Enumeration of immune cells from control SpiB^Flox/Flox^ and SpiB^ΔCCL19^ mice by flow cytometry. Graphs show pooled data (means ± SEM) from two independent experiments with 7-8 mice per group. **(B)** Skin draining lymph node sections from control SpiB^Flox/Flox^ and SpiB^ΔCCL19^ mice were stained for Lyve-1, CD31, PDPN, CD3 and B220 and analysed by confocal microscopy. Left images display the merged staining and right images the merged staining of CD3 and B220 only. Scale bar, 500 μm. **(C)** Quantification of the total lymph node size area (left graph) and lymph node compartment areas (right graph). Graphs show pooled data (means ± SEM) from two independent experiments with 5 mice per group. **(D-E)** TRC gap analysis on lymph node sections from control SpiB^Flox/Flox^ and SpiB^ΔCCL19^ mice. Sections were stained for PDPN (left) and converted to threshold images (middle) and overlayed with circles for gap analysis (right). Graph shows the quantification of circle radius. Each point represents a circle radius and data are pooled from 2 independent experiments with 5 mice per group. Scale bar, 50 μm. **(F)** Ratio analysis of total T cells versus TRCs in control SpiB^Flox/Flox^ and SpiB^ΔCCL19^ mice. Graph shows pooled data (means ± SEM) from two independent experiments with 8 mice per group *p < 0.05, ****p < 0.0001, ns, non-significant, by Mann-Whitney test (A, C, E and F).

### SpiB expression enables FRC to regulate T cell immunity

Having identified a role for SpiB in regulating the TRC function and network properties at homeostasis, we then investigated if SpiB expression in FRCs supports immune responses. For this, we labelled with CellTrace Violet (CTV) gBT-I CD8^+^ T cells specific for an Herpes simplex virus (HSV) glycoprotein B epitope and transferred into SpiB^ΔCCL19^ and control mice followed by subcutaneous HSV-1 KOS infection (Fig. 4A). We tracked CD45.1^+^ gBT-I CD8^+^ T cells in the draining popliteal LNs of infected mice (Fig. 4B). Proliferation of gBT-I CD8^+^ T cells 3 days after infection was diminished in SpiB^ΔCCL19^ mice, reflected by a lower average number of divisions (proliferation index) and reduced fold expansion (replication index) amongst dividing cells (Fig. 4B-C). Accumulation of divided gBT-I CD8^+^ T cells required SpiB expression in FRCs (Fig. 4D), yet upregulation of the activation markers CD69 and CD25 by gBT-I cells was unaffected, suggesting normal differentiation of the CD8^+^ T cells that entered division (Fig. 4E).

**Figure 4:**
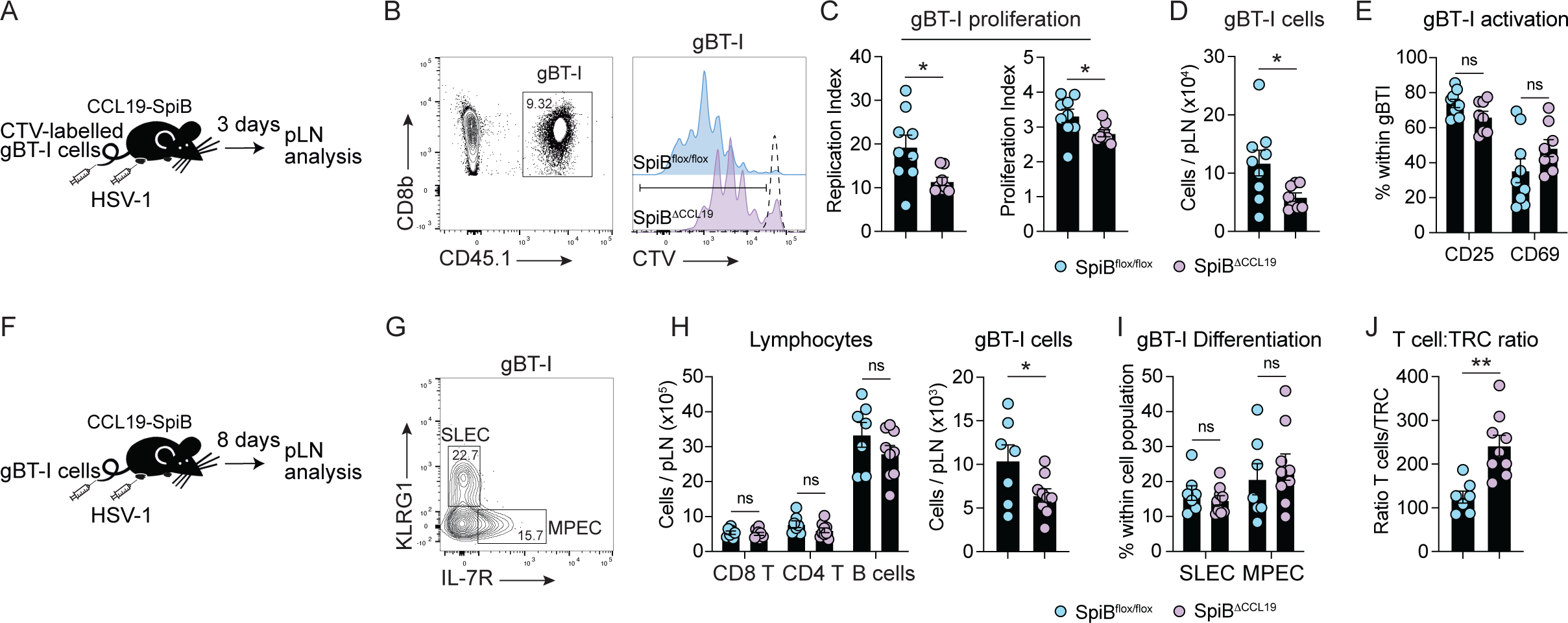
SpiB expression in FRC regulates T cell responses during viral infection. **(A)** Experimental schematic of HSV-1 infection for T cell priming. Mice were injected with 1.5×10^6^ CTV-labelled gBT-I cells and 24h later infected subcutaneously in the footpad with HSV-1. Draining popliteal LNs were analysed 3 days post-infection. **(B)** Flow cytometry analysis of CD45.1^+^ CD8^+^ gBT-I cells and representative histograms of CTV labelled gBT-I cells in the draining popliteal lymph node of control SpiB^Flox/Flox^ and SpiB^ΔCCL19^ mice, 3 days post subcutaneous HSV-1 infection. Dotted histogram represents CTV staining on gBT-I cells from naïve mice used to gate on divided gBT-I cells. **(C-E)** Quantification of gBT-I cell expansion and activation. (C) Left graph shows the replication index (calculated as Total Number of Divided Cells / Cells that Went into Division) and right graph the proliferation index (calculated as Total Number of Divisions / Cells that went into division) of gBT-I cells in the draining popliteal lymph node of control SpiB^Flox/Flox^ and SpiB^ΔCCL19^ mice. (D) Graph shows the enumeration of divided gBT-I cells, 3 days post subcutaneous HSV-1 infection. (E) Quantification of the percentage of CD25^+^ and CD69^+^ gBT-I cells from. Graphs show pooled data (means ± SEM) from two independent experiments with 9-8 mice per group. **(F)** Experimental schematic of HSV-1 infection. Mice were injected with 5×10^4^ gBT-I cells and 24h later infected subcutaneously in the footpad with HSV-1. Draining popliteal LNs were analysed 8 days post-infection. **(G-I)** Flow cytometry analysis of CD45.1+ gBT-I cells in the draining popliteal lymph node, 8 days post subcutaneous HSV-1 infection. (G) Short lived effector cells (SLEC) and memory precursor effector cells (MPEC) gBT-I cells were identified as KLRG1^+^ and IL-7R^+^ respectively. Enumeration of lymphocytes and gBT-I cells (H), and gBT-I cell differentiation (I) in the draining popliteal lymph node of control SpiB^Flox/Flox^ and SpiB^ΔCCL19^ mice, 8 days post subcutaneous HSV-1 infection. **(J)** Ratio analysis of total T cells versus TRCs in control SpiB^Flox/Flox^ and SpiB^ΔCCL19^ mice in the draining popliteal lymph node, 8 days post subcutaneous HSV-1 infection. Graphs (H-J) show pooled data (means ± SEM) from two independent experiments with 7-9 mice per group. *p < 0.05, **p < 0.01, ns, non-significant, by unpaired two-tailed t test (C-E) and Mann-Whitney test (H-J).

We then explored if the reduced early proliferation of gBT-I cells would further impact their effector functions. For this, we similarly tracked and analysed the differentiation of gBT-I cells into short lived effector cells (SLECs) and memory precursor effector cells (MPECs) based on expression of KLRG1 and the IL-7R respectively, in the draining popliteal LNs of infected mice 8 days post-infection (Fig. 4F-G). Although total numbers of CD8^+^ T, CD4^+^ T and B cells were similar in LNs of SpiB^ΔCCL19^ mice when compared to littermate control mice, we observed a significant reduction in numbers of virus-specific gBT-I CD8^+^ T cells (Fig. 4H). The differentiation of gBT-I cells into SLEC and MPEC was not affected (Fig. 4I), and induction of GL7^+^CD38^-^ germinal centre B cells, CD138^+^ antibody secreting cells and CXCR5^+^PD-I^+^ T_FH_ cells were not altered in the LNs of HSV-1 infected SpiB^ΔCCL19^ mice (Supp Fig. 2A-B). This suggested that SpiB expression in FRCs preferentially supported CD8^+^ T cell responses. To confirm this, we infected mice with Lymphocytic choriomeningitis virus and examined cellularity in inguinal LNs. Expansion of endogenous CD4^+^ and CD8^+^ T cells and virus-specific transgenic P14 CD8^+^ T cells was reduced in SpiB^ΔCCL19^ mice, but differentiation of effector CD8^+^ T cells was not altered (Supp Fig. 2C-D).

Finally, we observed a significant defect in the expansion of the TRC network during the course of HSV-1 infection in SpiB^ΔCCL19^ mice that resulted in a significant increase in the ratio of total T cells to TRCs in LNs (Fig. 4J, Supp Fig. 2E). These results suggest that SpiB expression in FRCs is critical for the expansion of the FRC network that regulates the early priming and proliferation of CD8^+^ T cells during viral infection. Together, our findings expand our understanding of the role of SpiB in regulating the LN FRC network and a crucial role in supporting the induction of CD8^+^ T cell responses.

## Discussion

This study identified the transcription SpiB as a regulator of FRC homeostasis and functionality in LNs. We found that in the absence of SpiB, FRCs were decreased in cellularity and key homeostatic factors expressed by FRCs such as CCL21 and IL-7 were reduced. However, we found that the size and architecture of LNs were not affected. In addition, we found that the absence of SpiB expression in FRCs did not affect the expression of canonical FRC markers such as PDPN, CD140a, BST1 and VCAM1, or the expression of other chemokines, including *Ccl19*, suggesting normal differentiation of FRCs from precursor cells. This is sharp in contrast with previous work that identified that the lymphotoxin beta signalling pathway or the expression of YAP/TAZ in CCL19^+^ FRCs are critical for the maturation of FRCs from precursor cells required for proper organisation of LN structures^18–20^. In these models, homeostatic chemokine expression such as CCL21 and CCL19 were also strongly reduced in FRCs that impacted the recruitment of immune cells leading to smaller LN size and deformed architecture, notably the delineation of the T and B cell compartments. We found that even though CCL21 expression was reduced in FRCs in the absence of SpiB, our histological examination of LNs showed normal CCL21 accumulation on the reticular network, suggesting that reduced chemokine expression in FRCs might be compensated from other sources such as endothelial cells, that are a source of the chemokine in lymphoid tissues^5,13^.

Despite normal compartmentalisation of the LN, we observed that the space between the reticular network fibers within the T cell zone of LNs was increased in the absence of SpiB expression. This implies that the FRC network has remodelled to maintain the size of the LN and ratio with T cells in the steady state. Under basal conditions, FRCs maintain tension via PDPN expression that facilitates the contraction of the actomyosin cytoskeleton^27,28^ and this contraction of the FRC network regulates the stiffness of the LN as well as their size. During the first few days of an immune response, CLEC-2 expression from mature dendritic cells engages and blocks PDPN functions and downstream signalling to relax the actomyosin cytoskeleton and induce the stretching of FRCs^27,28^. This regulates LN tension to accommodate the increase of cellularity due to lymphocyte recruitment^29–31^. We did not find that SpiB regulates PDPN expression in FRCs suggesting that additional molecules involved in FRCs contractility should be investigated in our model. A recent report identified that the PDPN binding partner surface proteins CD44 and CD9 suppress PDPN functions and the contractility of FRCs^32^. Whether SpiB regulates expression of both CD44 and CD9 remains to be addressed. In addition, the subcellular distribution of the transcription factors YAP and TAZ are a direct proxy of mechanical signalling that cells receive^33^ and two recent studies identified that the nuclear relocalisation of YAP/TAZ correlates with increased tension while their cytoplasmic localisation correlates with decreased tension and relaxation of the FRC network^30,34^. Our imaging indicates that the FRC network may be relaxed in the T cell zone of LNs and may indicate a lower nuclear/cytoplasmic ratio of YAP/TAZ in WT FRCs compared to their SpiB deficient counterparts.

Our data identified that SpiB expression in FRCs is required for the optimal activation of naïve CD8^+^ T cells during viral infection. In the absence of SpiB, the early proliferation of antiviral CD8^+^ T cells was delayed. This corroborates our previous findings where we identified a role for SpiB expression in spleen FRCs in regulating CD8 T cell responses to acute and chronic systemic viral infection^16^. Previous studies also identified a defect in CD8^+^ T cell responses when the FRC network was impaired in the absence of the lymphotoxin beta signalling pathway or the expression of YAP/TAZ in FRCs^18,19^. In these two studies, the decrease of FRC as well as reduction in homeostatic chemokine expression resulted in a paucity of immune cells in LNs, and disorganised T/B segregation that likely contributed to the reduced T cell response. Similarly, gradual depletion of the CCL19^+^ FRC network revealed that the topology of the network can accommodate up to 50% decrease before affecting immune cell recruitment to LNs, intranodal cell motility or priming of CD8^+^ T cells^35^. Given that we found only a small reduction in FRC numbers in SpiB^ΔCCL19^ LNs, our data therefore indicate that functional changes in FRC were responsible for defects in T cell priming. Yet, the mechanisms behind FRC support for T cell responses, beyond the chemokine-dependent positioning of immune cells, is still unclear. We identified that SpiB expression is required for LN remodelling via optimal FRC expansion during viral infection and that the ratio of T cells to FRC was increased over the course of infection suggesting that T cells would likely make less contacts with FRCs as the LN expands. Additionally, we cannot exclude the requirement for direct signals from FRCs for T cell activation that are regulated by SpiB but not identified in our study. Future studies would need to determine how SpiB regulates FRC functionality during inflammation to identify factors that could influence immune responses. In summary, our study establishes a role for the transcription factor SpiB in regulating LN FRC functions and provide new insights into how the FRC network support immune responses.

## Acknowledgements

We thank the Bioresources Facility and the Melbourne Cytometry Platform at the Peter Doherty Institute for technical support. Confocal Imaging was performed at the Biological Optical Microscopy Platform (BOMP) Facility at The University of Melbourne.

## Author contribution

Conceptualization: Y.O.A. and S.N.M. Methodology: H.L.H. and Y.O.A. Investigation: H.L.H. and Y.O.A. Writing (original draft): Y.O.A. Writing (review and editing): H.L.H., Y.O.A. and S.N.M. Resources: W.C. and G.T.B. Visualization: H.L.H. and Y.O.A. Supervision: Y.O.A and S.N.M. Funding acquisition: Y.O.A. and S.N.M.

## Material and methods

### Mice

C57BL/6, CCL19-Cre, SpiB^flox/flox^, gBT-I3B6.SJL-PtprcaPep3b/BoyJ (gBT-I.CD45.1), P143B6.SJL-PtprcaPep3b/BoyJ (P14.CD45.1) mice were bred in the Doherty Institute. SpiB-tdTomato were bred at WEHI. SpiB^ΔCCL19^ mice were generated by crossing CCL19-Cre and SpiB^flox/flox^ mice. Animal experiments were approved by the University of Melbourne Animal Ethics Committee. Mice were maintained under specific pathogen–free conditions and housed in individually ventilated cages. All mice were sex- and age-matched, and both female and male mice were used between 8 and 14 weeks of age.

### Adoptive transfer of transgenic CD8+ T cells

Recipient mice were intravenously injected with 5×10^4^ gBT-I or P14 cells 24 hours before infection. For priming experiments, gBT-I cells were first labelled with CellTrace Violet (ThermoFischer) at a final concentration of 5μM according to manufacturer’s instructions and 1.5 million cells were intravenously injected in recipient mice.

### Virus and infections

Herpes Simplex Virus-1 (HSV-1) KOS strain and Lymphocytic choriomeningitis virus Armstrong strain were used in this study. Mice were anaesthetized with a mixture of ketamine/xylazine at 100mg/kg and 20mg/kg respectively and infected subcutaneously in the footpad with 2 x 10^4^ of plaque-forming units (PFU) of HSV-1. Mice were infected intraperitoneally with 2 × 10^5^ PFU of LCMV Armstrong.

### LN digestion and Flow cytometry

LNs were incubated at 37°C in RPMI with 2mg/mL collagenase D, 0.8mg/mL Dispase and 100ug/mL DNase and 2% Fetal Bovine Serum. LNs were gently digested by removing and replacing the cell suspension every 10 min until completely digested. LN cell suspensions were resuspended in FACS buffer (PBS 2% BSA 5mM EDTA) and filtered through 70 μM before antibody staining. Cells were stained in FACS buffer containing CD16/32 Fc blocking antibody for 30min at 4°C. Antibodies used for staining are detailed in Table S1. Intracellular staining for CCL21 was performed using eBioscience Intracellular Fixation & Permeabilization Buffer Set according to the manufacturer’s instructions. Cells were enumerated by adding SPHERO calibration particles to each sample before acquisition on flow cytometer and samples were acquired using FACSFortessa (BD) or Cytek Aurora, and FlowJo software was used for analysis.

### Quantitative Real-Time PCR

Total RNA was extracted from sorted samples using RNeasy Plus Micro Kit (Qiagen) and converted to complementary DNA (cDNA) using the High Capacity cDNA Reverse Transcription Kit (Thermo Fisher Scientific) according to the manufacturer’s instructions. Genes of interest were preamplified from cDNA using TaqMan PreAmp Master Mix (Thermo Fisher Scientific) and samples were analysed by real-time qPCR using Fast SYBR Green Master Mix. Cycle-threshold values were determined for genes individually, and gene expression was normalized to the housekeeping genes Hprt and Gapdh (ΔCt) and presented as 2^−ΔCt^ [arbitrary units (AU)].

### Immunofluorescence and confocal imaging

LNs were harvested and fixed in 4% paraformaldehyde for 4 hours, incubated in 30% sucrose and embedded in OCT freezing media. Control and SpiB^ΔCCL19^ LNs were embedded in the same block for comparative analysis. Tissue sections were cut at 20μm thickness with a cryostat (Leica CM3050S). Sections were blocked for 2 hours (10% normal serum (NS), 0.3% Triton X-100 in PBS) at room temperature (RT). Sections were stained with primary antibodies (Table S2) overnight at 4 degrees (Diluted in PBS, 10% NS, 0.01% Triton X-100). Sections were washed in PBS-Tween 0.05% three times for 15 minutes and then blocked at room temperature for 2hrs. Secondary antibodies (Table S2) were applied for 2hrs at RT (Diluted in PBS, 10% NS, 0.01% Triton X-100). This was followed by two 15 minutes washes of PBS-Tween 0.05% and a final wash of 15 minutes in PBS. Sequential staining and blocking of CCL21 (goat) and PDPN (hamster) was performed to prevent cross reactivity. Stained sections were mounted in ProLong Gold antifade reagent and images acquired on a LSM980 confocal microscope (Carl Zeiss). Image analysis was performed in ImageJ. *Quantification of CCL21 and LN architecture.* Hand drawn ROIs of the LN were used to calculate the LN area. T, B cell zones and the medullary area were calculated by using the CD3, B220 and LYVE-1 positive stained region on maximum projections. T-cell zone FRCs regions of interest (ROI) were used to generate PDPN masks which were applied to CCL21 staining to calculate the fluorescent intensity within the fibroblast network. *Gap analysis*. The FRC gap analysis used a MATLAB script from the Acton lab^29^. PDPN fluorescence maximum projections were converted into a binary mask before a circle-fitting algorithm consecutively fit the largest circle possible within the gaps in the network that did not overlap with other fitted circles. Each circle was given a radius. The top 10 largest radii from each ROI were plotted.

### Statistical analysis

Graphs and statistics were generated using Prism 9 (GraphPad). Samples were tested for normality, and two groups were compared using two-tailed Mann-Whitney U test or unpaired t test. Multiple groups were analysed with one-way analysis of variance (ANOVA), followed by Tukey’s post-test comparison or Kruskal-Wallis, based on Gaussian distribution. All graphs depict means ± SEM. Details of statistical analysis are indicated in the figure legends and include the statistical test used. ns indicates nonsignificant; *P < 0.05, **P < 0.01, ***P < 0.001, and ****P < 0.0001.

---

**Supp Figure 1:**
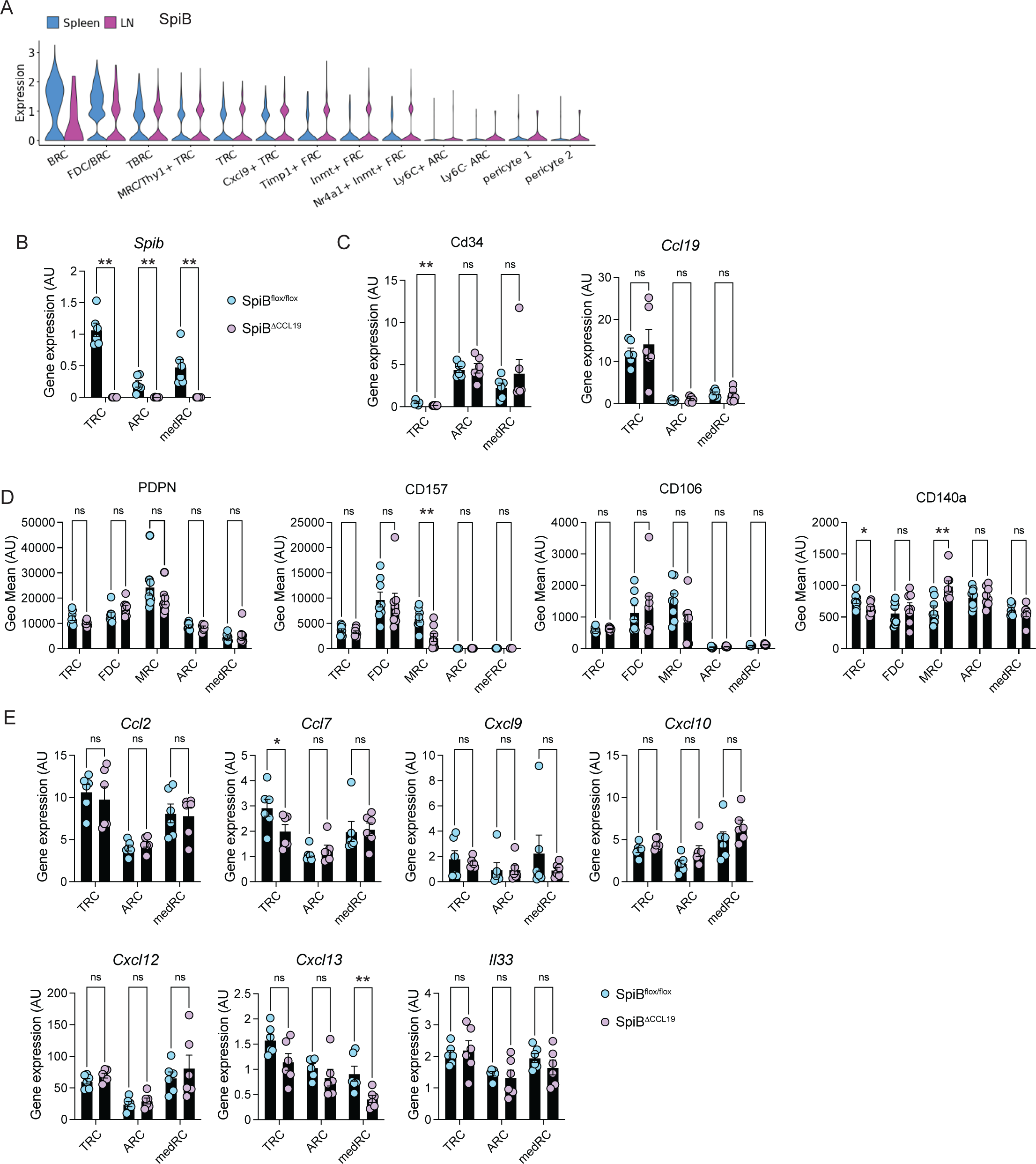
The transcription factor SpiB regulates TRC network and functionality. **(A)** Violin plots showing expression of *Spib* in Fibroblastic Reticular Cell subsets and pericytes in spleen and lymph node by single cell RNA sequencing. Data and plots were generated from http://muellerlab.mdhs.unimelb.edu.au/frc_scrnaseq/ . **(B-C)** Analysis of *Spib* (B), *Cd34* and *Ccl19* (C) expression in lymph node sorted TRCs, ARC and medRC of control SpiB^Flox/Flox^ and SpiB^ΔCCL19^ mice by qPCR. n = 6 mice from 3 independent sorts. **(D)** Flow cytometry analysis of PDPN, CD157, CD106 and CD140a expression in FRC subsets from control SpiB^Flox/Flox^ and SpiB^ΔCCL19^. Graphs display the Mean Fluorescence Intensity (GeoMean) of the different markers from pooled data (means ± SEM) from two independent experiments with 8 mice per group. **(E)** Analysis of *Ccl2, Ccl7, Cxcl9, Cxcl10, Cxcl12, Cxcl13* and *Il33* expression in lymph node sorted TRCs, ARC and medRC of control SpiB^Flox/Flox^ (WT) and SpiB^ΔCCL19^ (SpiB) mice by qPCR. n = 6 mice from 3 independent sorts. *p < 0.05, **p < 0.01, ns, non-significant, by unpaired two-tailed t test (D) and Mann-Whitney test (B, C and E).

**Supp Figure 2:**
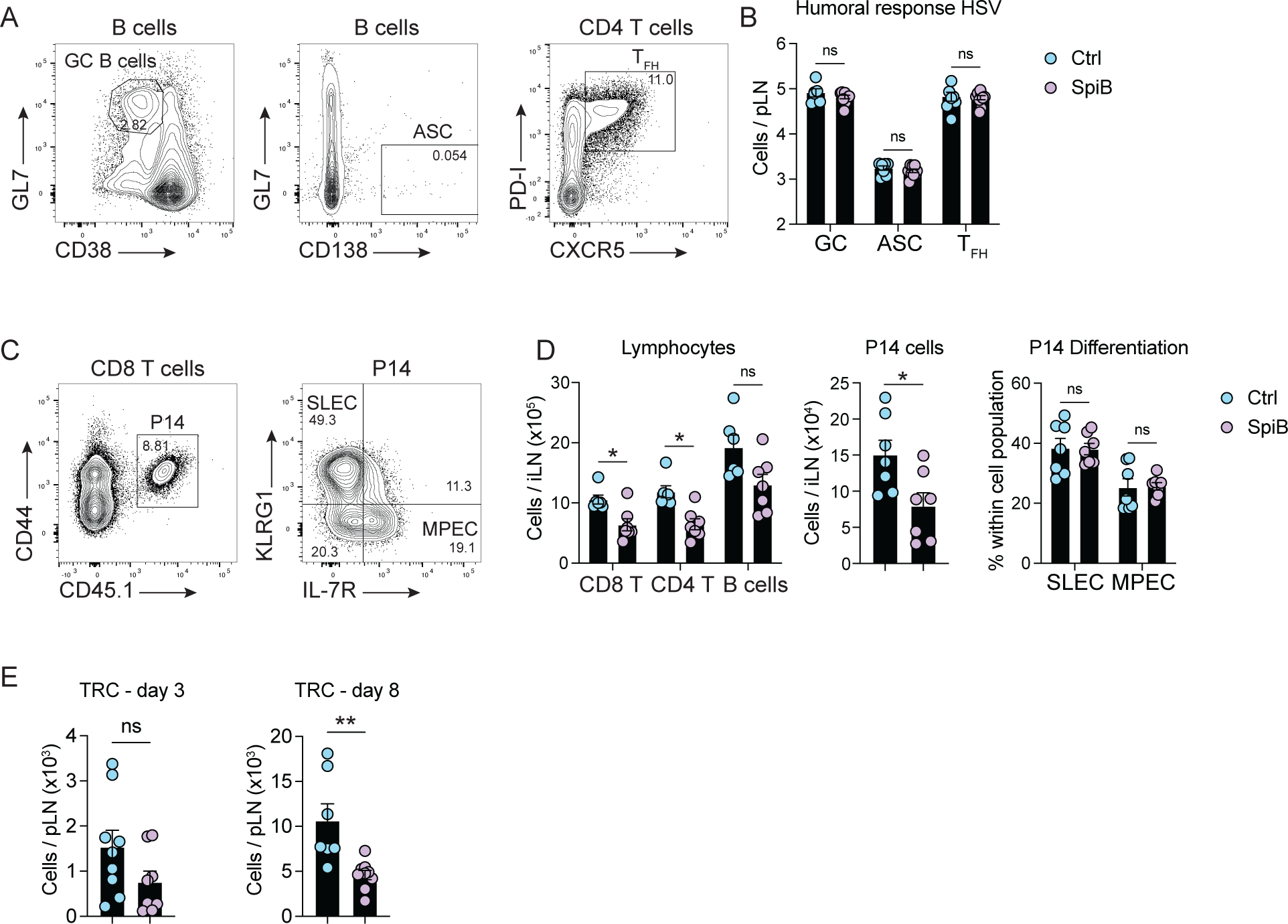
SpiB expression in FRC regulates T cell expansion during viral infection. **(A)** Flow cytometry analysis of Germinal centre (GC), antibody secreting (ASC) and T_FH_ cells in the draining popliteal lymph node of mice 8 days post-subcutaneous HSV-1 infection. **(B)** Enumeration GC, ASC and T_FH_ in the draining popliteal lymph node of control SpiB^Flox/Flox^ (WT) and SpiB^ΔCCL19^ (SpiB) mice, 8 days post subcutaneous HSV-1 infection. Graphs show pooled data (means ± SEM) from two independent experiments with 7-9 mice per group. **(C)** Flow cytometry analysis of CD45.1+ P14 cells in inguinal lymph node, 8 days post LCMV infection. Short lived effector cells (SLEC) and Memory precursor effector cells (MPEC) P14 cells were identified as KLRG1^+^ and IL-7R^+^ respectively. **(D)** Enumeration of lymphocytes, P14 cells, and P14 cell differentiation in inguinal lymph node of control SpiB^Flox/Flox^ (WT) and SpiB^ΔCCL19^ (SpiB) mice, 8 days post LCMV infection. **(E)** Enumeration of draining lymph node FRC from control SpiB^Flox/Flox^ (WT) and SpiB^ΔCCL19^ (SpiB) mice by flow cytometry at day 3 and 8 post-HSV-1 infection. Graphs show pooled data (means ± SEM) from two independent experiments with 9-8 mice (day 3) and 7-9 (day 8) mice per group. *p < 0.05, **p < 0.01, ns, non-significant, by Mann-Whitney test (B, D and E).

