## Supplementary material for "The transcription factor SpiB regulates Fibroblastic Reticular Cell network and CD8^+^ T cell responses in lymph nodes": Supp Table 1

| Supp Table | | | | | |
|---|---|---|---|---|---|
| Primary antibodies | | | | | |
| Name | Clone | Vendor | Catalog number | Usage | Dilution |
| PE-Cy7 anti-mouse Podoplanin | 8.1.1 | Life Technologies | 25-5381-82 | Flow | 1/500 |
| PerCP-Cy™5.5 anti-mouse CD21/CD35 | 7G6 | BD | 562797 | Flow | 1/100 |
| APC R700 anti-mouse CD45 | 30 F11 | BD | 565478 | Flow | 1/100 |
| Alexa Fluor® 700 anti-mouse TER-119/Erythroid Cells | TER-119 | BioLegend | 116220 | Flow | 1/200 |
| Brilliant Violet 421™ anti-mouse CD31 | 390 | BioLegend | 102424 | Flow | 1/500 |
| Brilliant Violet 605™ anti-mouse CD140a | APA5 | BioLegend | 135916 | Flow | 1/200 |
| Brilliant Violet 785™ anti-mouse Ly-6C | HK1.4 | Biolegend | 128041 | Flow | 1/500 |
| BV480 anti-mouse CD146 | ME-9F1 | BD | 746601 | Flow | 1/500 |
| BV650 anti-mouse CD157 | BP-3 | BD | 740611 | Flow | 1/500 |
| Alexa Fluor® 488 anti-mouse CD106 | 429 | Biolegend | 105710 | Flow | 1/200 |
| Biotin anti-mouse MAdCAM-1 | MECA-367 | Biolegend | 120706 | Flow | 1/100 |
| Purified anti-mouse Leptin R | Polyclonal | R&D Systems | AF497 | Flow | 1/200 |
| PE-Cy7 anti-mouse CD31 | MEC13.3 | Biolegend | 102418 | Flow | 1/500 |
| APC anti-mouse CD21/35 (CR2/CR1) | 7E9 | Biolegend | 123412 | Flow | 1/100 |
| BD Horizon™ APC-R700 Rat Anti-Mouse CD45 | 30-F11 | BD | 565478 | Flow | 1/200 |
| BD OptiBuild™ BUV563 Mouse Anti-Mouse NK-1.1 | PK136 | BD | 741233 | Flow | 1/200 |
| BD Horizon™ BUV737 Rat Anti-Mouse CD44 | IM7 | BD | 612799 | Flow | 1/500 |
| BD Horizon™ BUV805 Rat Anti-Mouse CD4 | RM4-5 | BD | 569193 | Flow | 1/200 |
| BD Horizon™ APC-R700 Mouse Anti-Mouse CD45.1 | A20 | BD | 565814 | Flow | 1/200 |
| BD OptiBuild™ BV750 Rat Anti-Mouse CD8b | H35-17.2 | BD | 747505 | Flow | 1/400 |
| BV480 Rat Anti-Mouse CD62L | MEL-14 | BD | 746726 | Flow | 1/500 |
| BV570 anti-mouse Ly-6C | HK1.4 | Biolegend | 128030 | Flow | 1/400 |
| BV650 anti-mouse CXCR3 | CXCR3-173 | Biolegend | 126531 | Flow | 1/200 |
| BV711 anti-mouse CD279 (PD-I) | 29F.1A12 | Biolegend | 135231 | Flow | 1/200 |
| BV785 anti-mouse CX3CR1 | SA011F11 | Biolegend | 149029 | Flow | 1/500 |
| FITC anti-mouse/human KLRG1 | 2F1 | Biolegend | 138410 | Flow | 1/200 |
| PE anti-mouse CD127 (IL-7Ra) | A7R34 | Biolegend | 121112 | Flow | 1/100 |
| PE-Cy™5 anti-mouse CD69 | H1.2F3 | Biolegend | 104510 | Flow | 1/400 |
| APC-Cy7 anti-mouse TCR beta chain | H57-597 | BD | 560656 | Flow | 1/200 |
| APC anti-mouse CD25 | 3C7 | Biolegend | 101910 | Flow | 1/200 |
| PE-Vio615 anti-mouse CD106 (VCAM-1) | REA971 | Miltenyi Biotec | 130116223 | Flow | 1/50 |
| eFluo450 anti-mouse CD90.2 (Thy1.2) | 53-2.1 | ThermoFischer | 48-0902-82 | Flow | 1/200 |
| BD Horizon™ BUV395 Rat Anti-Mouse CD45R/B220 | RA3-6B2 | BD | 563793 | Flow | 1/200 |
| BV421 anti-mouse F4/80 Antibody | BM8 | Biolegend | 123132 | Flow | 1/200 |
| V450 anti-mouse Ly-6G | 1A8 | BD | 560603 | Flow | 1/200 |
| Brilliant Violet 510™ anti-mouse/rat XCR1 | ZET | Biolegend | 148218 | Flow | 1/200 |
| Brilliant Violet 605™ anti-mouse CD11c Antibody | N418 | Biolegend | 117334 | Flow | 1/200 |
| Brilliant Violet 711™ anti-mouse/human CD11b | M1/70 | Biolegend | 101241 | Flow | 1/500 |
| PerCP/Cyanine5.5 anti-mouse CD19 Antibody | 6D5 | Biolegend | 115534 | Flow | 1/200 |
| PE anti-mouse CD192 (CCR2) | QA18A56 | Biolegend | 160106 | Flow | 1/200 |
| PeCy7 anti-mouse CD26 | DPP-4 | BioLegend | 137810 | Flow | 1/400 |
| APC anti-mouse CD64 (FcγRI) | X54-5/7.1 | BioLegend | 139306 | Flow | 1/200 |
| Alexa Fluor® 647 anti-mouse Siglec H | 551 | Biolegend | 129608 | Flow | 1/200 |
| Alexa Fluor™ 700 anti-mouse MHC Class II (I-A/I-E) | M5/114.15.2 | ThermoFischer | 56-5321-82 | Flow | 1/200 |
| Purified anti-mouse CCL21 | Polyclonal | R&D Systems | AF457 | Flow, IF | 1/100 |
| Purified anti-mouse Podoplanin | 8.1.1 | Biolegend | 127401 | IF | 1/100 |
| Pacific Blue™ anti-mouse/human CD45R/B220 | RA3-6B2 | BioLegend | 103227 | IF | 1/100 |
| Alexa Fluor® 594 anti-mouse CD3 Antibody | 17A2 | Biolegend | 100240 | IF | 1/100 |
| Alexa Fluor 647 anti Mouse CD31 | MEC13.3 | Biolegend | 102516 | IF | 1/100 |
| Purified anti-LYVE1 | Polyclonal | Abcam | ab14917 | IF | 1/500 |

| Secondary antibodies/reagents | | | | | |
|---|---|---|---|---|---|
| Name | Clone | Vendor | Catalog number | Usage | Dilution |
| Streptavidin Brilliant Violet 711™ | | BioLegend | 405241 | Flow | 1/200 |
| Streptavidin PE | | BD | 554061 | Flow | 1/1000 |
| Alexa Fluor™ 488 Donkey anti-Goat IgG (H+L) Cross-Adsorbed Secondary Antibody | Polyclonal | ThermoFischer | A-11055 | IF | 1/500 |
| Alexa Fluor™ 647 Goat anti-Syrian Hamster IgG (H+L) Cross-Adsorbed Secondary Antibody | Polyclonal | ThermoFischer | A-21451 | IF | 1/500 |
| Alexa Fluor™ 790 Goat anti-Rabbit IgG (H+L) Highly Cross-Adsorbed Secondary Antibody | Polyclonal | ThermoFischer | A11369 | IF | 1/500 |
